# A Bayesian approach for analysis of whole-genome bisulphite sequencing data identifies disease-associated changes in DNA methylation

**DOI:** 10.1101/041715

**Authors:** Owen J.L. Rackham, Sarah R. Langley, Thomas Oates, Eleni Vradi, Nathan Harmston, Prashant K. Srivastava, Jacques Behmoaras, Petros Dellaportas, Leonardo Bottolo, Enrico Petretto

## Abstract

DNA methylation is a key epigenetic modification involved in gene regulation whose contribution to disease susceptibility remains to be fully understood. Here, we present a novel Bayesian smoothing approach (called ABBA) to detect differentially methylated regions (DMRs) from whole-genome bisulphite sequencing (WGBS). We also show how this approach can be leveraged to identify disease-associated changes in DNA methylation, suggesting mechanisms through which these alterations might affect disease. From a data modeling perspective, ABBA has the distinctive feature of automatically adapting to different correlation structures in CpG methylation levels across the genome whilst taking into account the distance between CpG sites as a covariate. Our simulation study shows that ABBA has greater power to detect DMRs than existing methods, providing an accurate identification of DMRs in the large majority of simulated cases. To empirically demonstrate the method’s efficacy in generating biological hypotheses, we performed WGBS of primary macrophages derived from an experimental rat system of glomerulonephritis and used ABBA to identify >1,000 disease-associated DMRs. Investigation of these DMRs revealed differential DNA methylation localized to a 600bp region in the promoter of the *Ifitm3* gene. This was confirmed by ChIP-seq and RNA-seq analyses, showing differential transcription factor binding at the *Ifitm3* promoter by JunD (an established determinant of glomerulonephritis) and a consistent change in *Ifitm3* expression. Our ABBA analysis allowed us to propose a new role for *Ifitm3* in the pathogenesis of glomerulonephritis via a mechanism involving promoter hypermethylation that is associated with *Ifitm3* repression in the rat strain susceptible to glomerulonephritis.

## INTRODUCTION

One of the most important epigenetic modifications directly affecting DNA is methylation, where a methyl group is added to a cytosine base in the DNA sequence creating 5-methylcytosine. High-throughput sequencing techniques, such as whole-genome bisulphite sequencing (WGBS), now allow for genome-wide methylome data to be collected at single base-resolution (Harris *et al.* 2010). However, the challenge remains on how to accurately identify DNA methylation changes at the genome-wide level and also account for the complex correlation structures present in the data. Whilst it is still not fully understood how DNA methylation affects gene expression, it has been shown that depending on the location of the modification it can either have a positive or negative effect on the level of expression of genes (Gutierrez-Arcelus *et al.* 2013). How methylation patterns are regulated is complex and a full understanding of this process requires elucidating the mechanisms for *de novo* DNA methylation and demethylation, as well as the maintenance of methylation (Chen and Riggs 2011) However, the majority of functional methylation changes are found in methylation sites where cytosines are immediately followed by guanines, known as CpG dinucleotides (Ziller *et al.* 2011). These are not positioned randomly across the genome but tend to appear in clusters called CpG islands (CpGI) (Deaton and Bird 2011). It has been also shown that there are concordant methylation changes within CpGI and in the genomic regions immediately surrounding CpGI (also known as CpGI shores or CpGS). These “spatially correlated” DNA methylation patterns tend to be more strongly associated with gene expression changes than the methylation changes occurring in other parts of the genome (Gutierrez-Arcelus *et al.* 2015). The correlation of methylation levels between CpG sites is also highly dependent on their genomic context, varying greatly depending on where in the genome they are located (Zhang *et al.* 2015). For computational convenience, the dependence of methylation patterns between CpG sites is sometimes ignored by methods for differential methylation analysis. Alternatively, a simplified estimation of the correlation of methylation levels between neighbouring CpG sites (Bell *et al.* 2011) based on a user-defined parameterization of the degree of smoothing is introduced. These strategies might not be appropriate across different experimental scenarios and instead we propose an automatic probabilistic smoothing procedure of the average methylation levels across replicates (hereafter methylation profiles).

Beyond the initial univariate analysis of methylation changes at each individual CpG (for instance, using the Fisher’s exact test), recently the focus has shifted to identifying differentially methylated regions (DMRs), since coordinated changes in CpG methylation across genomic regions are known to impart the strongest regulatory influence. To this aim, a number of tools have been proposed to detect DMRs from WGBS data. Typically, these methods normally take one of two approaches: Either model the number of methylated/unmethylated reads using a binomial, negative-binomial distribution or discrete distributions with an over-dispersion parameter) such as MethylKit (Akalin *et al.* 2012), MethylSig (Park *et al.* 2014) and DSS (Feng *et al.* 2014). Alternatively in order to account for the correlation of methylation profiles between neighbouring CpG sites, a smoothing operator is applied in tools like BSmooth (Hansen *et al.* 2012), BiSeq (Hebestreit *et al.* 2013), DSS-single (Wu *et al.* 2015) – reviewed in Robinson *et al.* 2014 and Yu and Sun 2016b). Methods based on spline- (Hansen *et al.* 2012), kernel- (Hebestreit *et al.* 2013) perform generally well in practical applications. However, their results and the identification of the DMRs depend on the choice of the smoothing parameters values, e.g., window size or kernel bandwidth, a feature that makes them less general and prone to perform unequally when the default parameters values are changed. In these cases, smoothing parameters tuned by time-consuming sensitivity analysis based on different parameterizations is usually recommended, although this strategy is rarely applied in real data analyses. Other approaches, e.g., metilene (Jühling *et al.* 2015), propose segmentation algorithms to detect DMRs between single/groups of replicates without making any model assumption about the data generating mechanism and less dependent on the parameters definition. Furthermore, several other algorithms have been introduced, e.g., MOABS (Sun *et al.* 2014), Lux (Äijö *et al.* 2016), and MACAU (Lea *et al.* 2015), showing that bisulfite sequencing data analysis is an active area of research.

To address this dependence on parameterization and the subsequent lack of generality, we propose a fully Bayesian approach, approximate Bayesian bisulphite sequencing analysis (or ABBA), designed to smooth automatically the underlying - not directly observable – methylation profiles and reliably identify DMRs whilst borrowing information vertically across biological replicates and horizontally across correlated CpGs (**Fig. 1**). We highlight that this fully Bayesian specification is not adopted by previous DMR detection techniques, owing to the computational overhead of the inferential procedure. We address the high computational demands by utilizing a highly efficient inferential tool (Rue *et al.* 2009) for Bayesian models (see below and **Methods**). To demonstrate the benefits of adopting ABBA over existing approaches, we report a comprehensive simulation study where we benchmarked ABBA against five commonly used alternative methods (Fisher Exact Test, BSmooth, MethylKit, MethylSig, DSS) and considered a proposed new one (metilene) and assessed the effect of a different biological and experimental conditions (by varying parameters related to data integrity and quality of the signal) on the performance of each method. The results from this benchmark clearly indicate that ABBA is the best performing method, being both robust to changes in factors affecting data quality (e.g., sequencing coverage, errors associated with the methylation call) and level of noise in methylation signal. To benchmark our proposed method on a real dataset, we generated new WGBS data in macrophages from an established rat model of glomerulonephritis (Aitman *et al.* 2006) and control strain, and used ABBA for the genome-wide identification of DMRs. An additional comparison performed with the best alternative method (that arose from the simulation study) showed that ABBA has increased power to detect changes in DNA methylation involving genes and pathways relevant to glomerulonephritis. Furthermore, this comparison exemplifies how the DMR results obtained by alternative approaches depend heavily on the choice of relevant smoothing parameters (e.g., window size used in DSS). We also integrated the DMR results of ABBA with transcription factor binding site analysis, RNA-seq and ChIP-seq data generated in the same system, and in this we revealed a previously unappreciated role for the *Ifitm3* gene in the pathogenesis of glomerulonephritis, providing a proof of concept for real data applications of the ABBA approach.

**Figure 1.**
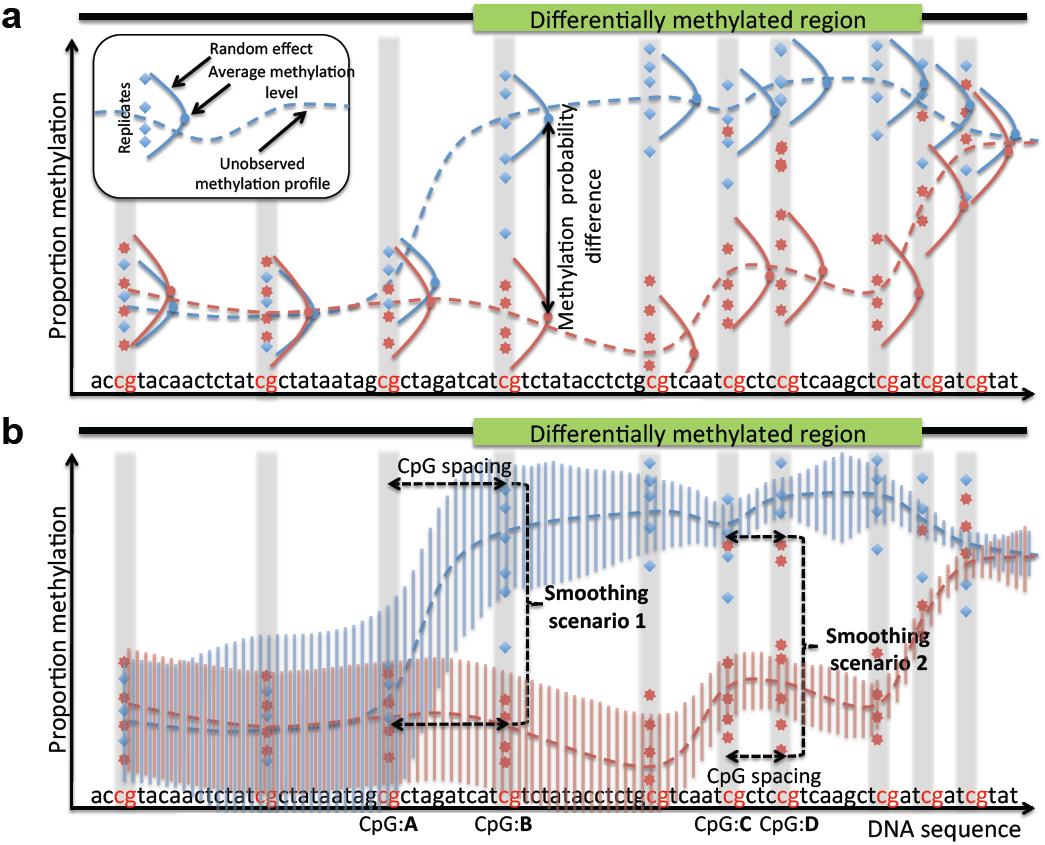
ABBA model. ABBA estimates the unobserved methylation profiles, i.e. the DNA average methylation levels across replicates, of two groups from WGBS data (blue diamonds and read stars). (**a**) A random effect accounts for the variability of experimental replicates. At each CpG the methylation probability difference is the difference between the methylation profile of the two groups (blue and red dots). (**b**) The methylation profiles of each group are smoothed by a latent Gaussian field that probabilistically connects them (dotted lines). In particular “Smoothing scenario 1” shows that if a large spacing (distance) between two consecutive CpGs (CpG:**A** and CpG:**B**) exists, the methylation profile at CpG:**B** does not depend on the previous one at CpG:**A** (blue dotted line). The opposite happens in “Smoothing scenario 2” where the methylation profile at CpG:**D** is largely influenced by the previous one at CpG:**C** (red dotted line) despite some high levels of methylation (red stars) which are treated by ABBA as outliers. The degree of the smoothing, i.e. the correlation between DNA methylation profiles, is controlled automatically by the marginal variance of the Latent Gaussian Field (blue and red vertical bars): the correlation is higher (lower) when the variance is small (large). On the other hand, the variance decreases as the distance between neighbouring CpGs’ decreases (Smoothing scenario 2) while increases as the distance increases (Smoothing scenario 1).

## MATERIALS AND METHODS

Below we reported the key aspects of the latent Gaussian model and Integrated Nested Laplace Approximation (INLA). Interested reader can also refer to Rue and Martino 2007 and Rue *et al.* 2009.

### Latent Gaussian model

A latent Gaussian model (LGM) can be described by a three-stage hierarchical model

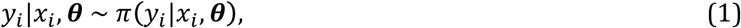

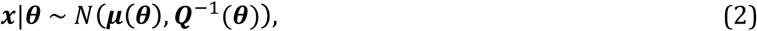

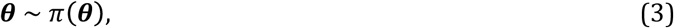

where *y*_*i*_ = 1,⋯, *n*, are the observed values, ***x*** is *n*-dimensional vector of latent variables and ***θ*** is *p*-dimensional vector of model parameters. (1) is the *observations equation* and it describes the probabilistic model for each observation conditionally on the latent variable *x*_*i*_ and the model parameters ***θ***, (2) is the *latent Gaussian field equation* with the latent variables distributed as a *p*-dimensional normal distribution, with mean vector ***μ***(***θ***) and a sparse precision matrix ***Q***(***θ***). Both quantities can depend on the model parameters vector ***θ*** whose distribution is described in the *parameter equation* (3). The Gaussian vector ***x*** exhibits a particular conditional dependence (or Markov) structure which is reflected in its precision matrix ***Q***(***θ***).

### Integrated Nested Laplace Approximation

INLA is a computational approach to perform statistical inference for LGM. It provides a fast and accurate alternative to exact MCMC (Gilks *et al.* 1996) and other sampling-based methods such as Sequential Monte Carlo (Doucet *et al.* 2001). They become prohibitively computationally expensive when the length of the sequence considered is too long, resulting in infeasible run times. The INLA solution with a mix of Laplace approximations (Tierney and Kadane 2012) and numerical integrations offers a pragmatic inferential tool to fit LGMs and it provides answers in hours whereas MCMC requires days. The INLA inferential procedure consists of three steps:

1. Compute the approximation to the marginal posterior
 *π*(***θ***|**y**) and by-product to *π*(*θ*_*j*_|***y***), *j* = 1,⋯,*p*;
2. Compute the approximation to *π*(*x*_*i*_|***y, θ***), *i* = 1,⋯, *n*;
3. Combine 1 and 2 above and compute the approximation to the marginal posterior *π*(*x*_*i*_|***y***).

### ABBA model

Based on LGM, the ABBA model can be described by a three-stage hierarchical model:

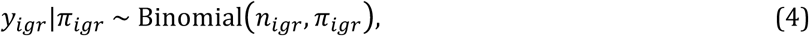

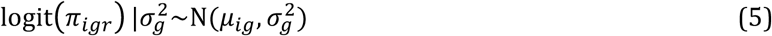

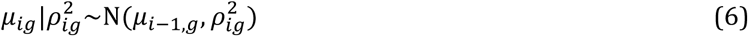

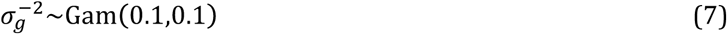

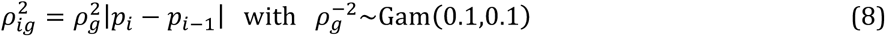

(4) is the *first part of the observations equation* where *i* = 1,⋯, *m* denotes the CpG, *g* = 1,2 the group (e.g., case and control group), and *r* = 1,⋯, *R* the experimental replicate. *y*_*igr*_, *n*_*igr*_ and *π*_*igr*_ are the observed number of methylated reads, the read depth and the proportion of methylation for the *i*th CpG site, *g*th group and *r*th experimental replicate, respectively. (5) is the *second part of the observations equation* and it describes a random effect across the experimental replicates with a specific variance 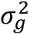 for each group. In (5), logit(*z*) indicates the logit transformation, logit(*z*) = log (1/(1 − *z*)). The observation equation (5) assumes that the methylation proportions are drawn from the same distribution within each group but are different between groups.

(6) is the *latent Gaussian field (LGF) equation.* The dependence of the DNA methylation pattern between CpGs is modelled as a non-stationary random walk of order 1, RW(1): *μ*_*ig*_ follows a normal distribution with mean *μ*_*i*−1,*g*_ (defined in the (*i* − 1)th CpG) and variance 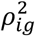 which is specific for each CpG and group. (5) and (6) highlight an important feature of ABBA model that is able to model vertically the information contained in the replicates by a random effect model and horizontally the information about the CpG methylation levels correlation by a LGF.

The model is completed by specification in (7) and (8) of the random effect and LGM prior precision, i.e. the inverse of the variance. For computational convenience we introduce a CpG site spacing and decompose 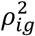 into 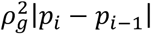, where 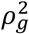 is the global smoothing parameter specific for each group that needs to be estimated and *p*_*i*_ and *p*_*i*−1_ are the chromosomal locations of two consecutive CpG sites. This implies that the correlation between *μ*_*ig*_ and *μ*_*i*−1,*g*_, depends on the distance between the two consecutive CpG sites and in particular, it decreases as this distance increases in keeping with empirical evidence reported in Bell JT et al. 2011, Zhang (2015) and in our real data set (see **Supplementary Figure 1**). With this formulation only 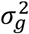 and 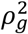 need to be estimated for each group. It also implies a sparse precision matrix ***Q***(***θ***) for the LGF in (2) making the overall inferential process efficient.

Finally, non-informative priors are assigned to the precision parameters 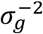 and 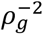 which are distributed as a gamma density with mean 1 and variance 10 (default INLA values). Sensitivity analysis on the gamma density parameterization shows no departure from the results obtained using the default values. See **Supplementary Table 1** for details on the posterior density of 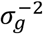 and 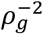 under INLA default and alternative parameterization on selected simulated examples.

When a single replicate is available, since 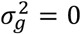, (4) and (5) simplify to

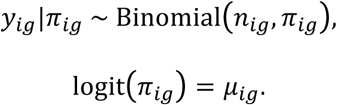

While some methods for DMR detection (Feng *et al.* 2014; Wu *et al.* 2015), allow for over-dispersion by assuming a beta-binomial model, (4) and (5) imply a logistic-normal model. After integrating out (6), 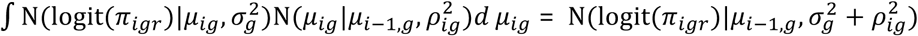, it can be shown that marginally

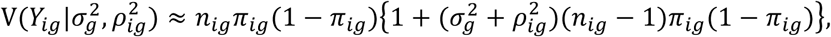

where *π*_ig_ ≡ exp(*μ*_*ig*_)/(1 + exp (*μ*_*ig*_)). The above equation illustrates that *a priori* the marginal degree of variability per CpG site under ABBA model is the variance of the binomial model multiplied by an over-dispersion factor that depends on the combined effect of 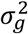, the replicates variability, and 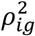, the variance of the unobserved methylation profile. When a single replicate is available, the over-dispersion depends only on 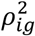.

#### ABBA algorithm

The ABBA algorithm consists of two steps:

1. Compute the approximation to the marginal posteriors of 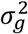, the variance of the random effect, and 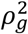, *g* = 1,2 the smoothing parameters; given the model specification 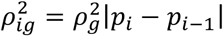, it is also possible to derive the marginal posteriors of 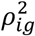;
2. Compute the approximation to marginal posterior *π*(*μ*_*ig*_|***y***), where ***y*** = (*y*_*igr*_)_*i*=1,⋯,*R*_; then the marginal posterior of the unobserved methylation profile *π*(*π*_*ig*_|***y***) is obtained by using the inverse logit transformation of *μ*_*ig*_, *z* ≡ exp{logit(*z*)}/[1 + exp{logit(*z*)}].

### Global differential methylation and FDR calculation

ABBA inference about DMRs is based on the posterior methylation probability (PMP) *π*(*π*_*ig*_|***y***) and the posterior differential methylation probability (PDMP) *π*(*π*_*i1*_|***y***) − *π*(*π*_*i2*_|***y***). The posterior mean methylation probability E(*π*_*ig*_|***y***) summarizes the information contained in the PMP and it is used to define the posterior mean differential methylation between two groups, *d*_*i*_ = E(*π*_*i1*_|***y***) − E(*π*_*i2*_|***y***). Once the LGF has been integrated out by INLA inferential process, *π*(*π*_*ig*_|***y***), *i* = 1,⋯, *n*, and in turn *d*_*i*_ s become marginally independent. This allows the straightforward application of a non-parametric false discovery rate (FDR) procedure without the burden of correlated signals. To distinguish between the null distribution (no differential methylation) and the alternatives, we fit a mixture of three truncated normal densities

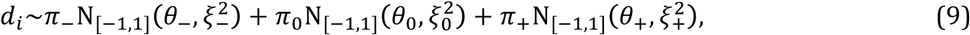

where N_[−1,1]_ is a normal density truncated between [−1,1], *π*_−_, *π*_0_, *π*_+_ ∈ (0,1) with *π*_−_ + *π*_0_ + *π*_+_ = 1 are the mixing weights of the “negative” differentially methylated, no differentially methylated and “positive” differentially methylated with respect the control group, respectively, *θ*_−_, *θ*_0_, *θ*_+_ are the unknown centers of the differentially methylated groups and 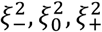 are the unknown variances. Under the null hypothesis we set *θ*_0_ = 0. For identifying the components of mixture model we also impose the condition *π*_0_ ≥ *π*_−_ + *π*_+_ under the assumption that the large majority of CpG sites are not differentially methylated.

Although the choice of a three component mixture model works well in real data examples (see **Supplementary Figure 2**), this assumption can be relaxed. For instance, as suggested in Sun and Cai 2009, the non-null distribution *f*_1_ can have more than two components. This allows a better fitting of the tails of distribution of *d*_*i*_’s and the identification of more than two differentially methylated groups. For instance the choice of the number of components can be based on Bayesian Information Criterion (BIC). However this requires running the FDR procedure several times for each choice of the number of components. Another possibility which is less computational intensive relies on the approximation of *f*_1_ by using a non-parametric Gaussian kernel density estimation (Kuan and Chiang 2012).

Maximum likelihood estimates of (9) are obtained by the EM algorithm (Dempster *et al.* 1977) taking particular care to avoid local maxima in the likelihood surface by running the EM algorithm from different starting points. Using the EM algorithm, the posterior probability of a CpG site belonging to each of the three component is

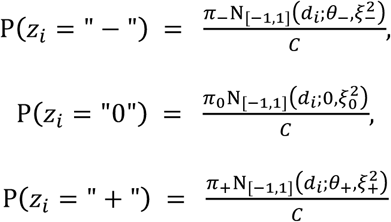

with 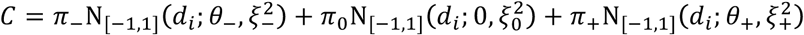.

Similarly to Broët *et al.* 2004, for a constant *t*, we define the estimated FDR(*t*) as

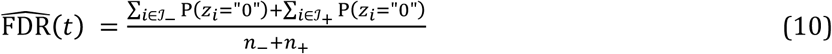

where 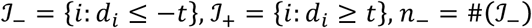 and 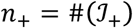. (10) defines the global FDR as the average local FDR which, for posterior probabilities, is defined as 1 − P(*z*_*i*_ = " − ") − P(*z*_*i*_ = " + " = P(*z*_*i*_ = "0"). Finally the constant *t* is chosen such that 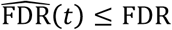.

In summary, the FDR procedure for ABBA consists of two steps:

1. Fit a mixture of truncated normal densities with three components on the *d*_*i*_s values; obtain the posterior probability that each *d*_*i*_ belongs to each of the three components;
2. Calculate the constant *t* such that 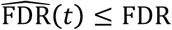 for a desired level of FDR;

For computational efficiency our FDR procedure can be run on each chromosome separately and then the results can be aggregated at the genome-wide level (Efron 2008). Besides the computational speed, this strategy does not assume the existence of a global methylation level difference between the two conditions that may not hold in practice. The separate-class model (Efron 2008), can be used to combine separate chromosome-wide FDRs.

### WGBS data simulation

WGBS data have a number of intrinsic characteristics that can vary depending on the cell-types/tissue complexity being studied or on technical issues related to the sequencing. In order assess which method is the most robust for analyzing WGBS data it is important that changes in each of these characteristics are taken into account. Here we take advantage of our previously published WGBS-data simulator (Rackham *et al.* 2015) that allows us to generate unbiased benchmarking datasets with several varying parameters. Wherever possible we will refer to the notation used in Rackham *et al.* 2015; the parameters are the following:

1. Number of replicates - the parameter *r* was set to vary between *r* = 1,2,3 within each group;
2. Average read depth – at each CpG site for all replicates and groups, the number of reads *n*_*igr*_, *i* = 1,⋯, *m* and *g* = 1,2, is simulated using a Poisson distribution with average read depth *λ*. The parameter *λ* was set to be either 10 or 30 reads on average per CpG site;
3. Level of noise - the parameter *s*_0_ controls the level of noise added the probability of methylation at each CpG site for all replicates and groups and simulates the measurement error resulting from the sampling of DNA segments during sequencing

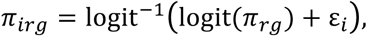

where *π*_*rg*_ is the global probability of methylation of the binomial (emission) distribution based on the real dataset analyzed (see details in Rackham *et al.* 2015) and ε_*i*_~N(0, *s*_0_), *i* = 1,⋯, *m*. *s*_0_ was set to vary between 0.1, 0.2 and 0.3 to model different level of noise. To calibrate the value of *s*_0_, **Supplementary Table 2** provides a Monte Carlo estimation of the effect of different values of the noise level on *π*_*irg*_.
4. Methylation probability difference - the parameter *Δmeth* reported in Rackham *et al.* 2015 as “phase difference” controls the magnitude of the difference between the probabilities of methylation in each group and was set to vary between 20%, 30%, 50% or 70%. This difference is obtained on CpG sites where both case and control samples share the same methylated status (methylated or unmethylated), by adding a given value to the probability in either cases or controls. The total length of the sequence where this difference appears in no greater than 5% (WGBSSuite default value) of the total length of the simulated region.
5. We also considered an additional parameter *δ* (not available for modeling in WGBSSuite), which introduces a further error associated with the methylation call. After selecting at random with a given probability *δ* a CpG site in the *g*th group for all replicates, we switch its methylation status between the two groups. In our simulation study, the parameter *δ* has been varied from 0, 0.05 and 0.1.

To perform the benchmarking we generate 5 replicates of 5,000 CpGs for each combination of the above parameters. The resulted in a total of 216 benchmarking datasets (3 cases for the number of replicates, 2 cases for the average read depth, 3 cases for the level of noise, 4 cases for the methylation probability difference, 3 cases for the parameter *δ*) which are replicated 5 times (5,400,000 CpGs in total) to assess the Monte Carlo average performance for each combination of parameters. In these datasets the size of the differentially methylated regions has a median size of 15 CpGs (see **Supplementary Figure 3**). The proportion of differentially methylated CpGs cannot exceed 20% of all CpGs (i.e., ~1000 CpGs).

### Receiver operator curve (ROC) construction for benchmarking

In order to generate the ROC curve the performance is calculated CpG-wise. For a given DMR, detection of each of the CpG contained within is considered as a true positive, whilst CpGs that are not detected are considered false negatives. Outside of the DMR the opposite criteria is applied. We choose this assignment criteria rather than calling detection of a each DMR since it provides a useful quantification of the extent each DMR is captured by each technique, for instance if one technique correctly identifies all the CpGs in a DMR, the method is deemed to perform better than an approach that identifies correctly only 80% of the CpGs within the same DMR.

### WGBS data pre-processing for ABBA

To run ABBA efficiently at the genome-wide level we took advantage of cluster-computing environment that enables parallel computation, and to this aim we preprocessed the WGBS data as follows. After the raw WGBS data were aligned, we removed CpG sites where less than 50% of the samples contain reads. Next, we split the WGBS data into chunks such that the distance between the last CpG site in one chunk and the first CpG in the next chunk is greater than 3,000bp. It has been previously shown that the correlation of DNA methylation levels between CpG sites decreases dramatically after 400bp (Zhang *et al.* 2015), so splitting the data in this way implies a particular conditional dependence structure in our data defined by a sparse block-diagonal precision matrix ***Q***(***θ***) where each block corresponds to a WGBS chunk. Chunks are then analyzed in parallel in a cluster-computing environment. We calculated the time required by ABBA to analyse chunks of different length (that span from 100 CpGs to 15,000 CpGs) on a single machine with 20 2.3GHz hyper-threaded cores and 32GB of RAM and found that the computational time (seconds) scales with the chunk length (*N*_CpG_, number of CpG sites) following the power function: *time* (seconds) = 0.0045 *N*_CpG_ ^1.3985^ (*R*^2^ = 0.997). Depending on the genome length and data dimensionality a complete WGBS analysis ABBA might require days (e.g., it took ~2 weeks to analyse WGBS data in the rat). The total computational time of ABBA analysis can be significantly shortened by splitting the genome into smaller chunks and then assemble the result. The results provided by the “whole-genome” ABBA analysis and “smaller-chunks” ABBA analyses are highly consistent, with no differences in the distribution probabilities obtained with and without splitting the genome in chunks (**Supplementary Figure 4**). Scripts for the pre-processing step are embedded within ABBA at abba.systems-genetics.net

### WGBS of rat macrophages

Bone-marrow derived macrophages (BMDM) were isolated from WKY and LEW rat strains. WGBS libraries were produced as follows: 6μg of genomic DNA was spiked with 10ng of unmethylated cl857 Sam7 lambda DNA (Promega) and sheared using a Covaris System S-series model S2. Sheared DNA was purified and then end-repaired in a 100μl reaction using NEBNext End Repair kit (New England Biolabs) incubated at 20C for 30 minutes. End-repaired DNA was next A-tailed using NEBNext dA-tailing reaction buffer and Klenow Fragment (also New England Biolabs) incubated at 37C for 30 minutes and then purified with the MinElute PCR purification kit (Qiagen) in a total final elution volume of 28μl. Illumina Early Access Methylation adapter oligos (Illumina) were then ligated to a total of 25μl of the A-tailed DNA sample using NEBNext Quick Ligation Reaction Buffer and Quick T4 DNA ligase (both New England Biolabs) in a reaction volume of 50μl. This mixture was incubated for 30 minutes at 20C prior to gel purification. Bisulphite conversion of 450ng of the purified DNA library was achieved using the Epitect Bisulfite kit (Qiagen) in a total volume of 140μl. Samples were incubated with the following program: 95C for 5 minutes, 60C for 25 minutes, 95C for 5 minutes, 60C for 85 minutes, 95C for 5 minutes, 60C for 175 minutes and then 3x repeat of 95C for 5 minutes and 60C for 180 minutes and held at 20C. Treated samples were then purified as per manufacturers instructions. Adapter bound DNA fragments were amplified by a 10-cycle PCR reaction and then purified using Agencourt AMPure XP beads (Beckman Coulter) before gel extraction and quantification using the Agilent Bioanalyzer 2100 Expert High Sensitivity DNA Assay. Then, libraries were quantified using quantitative PCR and then denatured into single stranded fragments. These fragments were then amplified by the Illumina cluster robot and transferred to the HiSeq 2000 for sequencing. WGBS reads were aligned and filtered according to a previously published pipeline (see (Johnson *et al.* 2012) and (Johnson *et al.* 2014)). Briefly, reads were pre-processed by in silico conversion of C bases to T bases in read 1 and G bases to A bases in read 2, followed by clipping of the first base from each read. Pre-processed reads were aligned to the rat reference genome (RGSC3.4) using BWA version 0.6.1 (Li and Durbin 2009) with 3’ end quality trimming using a Q score cutoff of 20. Converted and clipped reads 1 and 2 were mapped to two in silico converted versions of the reference sequence, firstly with Cs converted to Ts to allow forward strand mapping, and secondly with Gs converted to As to allow mapping of reverse strand. Aligned reads were filtered by removal of clonal reads, reads with a mapping quality of <20, reads that mapped to both in silico converted forward and reverse strands, and reads with an invalid mapping orientation. We obtained 79.9 billion ‘mappable’ bases across both rat strains, with 13.5x (average) coverage in the Lew strain and 17.6x (average) in WKY, where the greatest depth of coverage was observed within CpG islands.

Despite ABBA being able to detect methylation changes at all genomic locations we focused only on those methylation changes that occur at CpG sites, and considered CpG sites where at least 4 out of the 8 samples contain reads (resulting in a total of 14,976,632 CpG sites genome-wide in BMDM from WKY and LEW rats). DMRs were called with ABBA (see above) using a 5 CpG minimum, a 33% or greater difference in methylation and a 5% FDR threshold. Genomic region annotations and Ensembl gene IDs for the rat reference genome 4 (rn4) were downloaded from the UCSC genome browser. Significant over-representations of genomic features (intron, exons, etc.) were determined empirically from 1,000 randomly sampled length and GC-matched regions per DMR. The genes overlapping with DMRs were further annotated and tested for enrichment in Kyoto Encyclopedia of Genes and Genomes (KEGG) pathways using WebGestalt (Wang *et al.* 2013).

Identification of enriched transcription factor binding site (TFBS) motifs within the DMRs identified by ABBA was performed using HOMER (Heinz *et al.* 2010). HOMER was used to scan for motifs obtained from the JASPAR 2014 database (Mathelier *et al.* 2014). Threshold used for motifs identification was a p-value of 10^-4^. Enrichments were calculated by comparing the motifs present in the DMRs against a large set of background sequences (*N* = 10^6^) corrected for CpG content.

### RNA-seq and ChIP-seq analysis of rat macrophages

RNA-seq data from BMDM in WKY and LEW strains were retrieved from (Rotival *et al.* 2015) and reanalyzed in the context of WGBS analysis reported here. Briefly, total RNA was extracted from BMDM at day 5 of differentiation in three WKY rats and three LEW rats using Trizol (Invitrogen). 1 μg of total RNA was used to generate RNA-seq libraries using TruSeq RNA sample preparation kit (Illumina, UK). Libraries were run on a single lane per sample of the HiSeq 2000 platform (Illumina) to generate 100bp paired-end reads. An average depth of 72M reads per sample was achieved (minimum 38 M). RNA-seq reads were aligned to the rn4 reference genome using tophat2. The average number of mapped was 67M (minimum 36M) corresponding to an average mapping percentage of 93%. Sequencing and mapping were quality controlled using the FastQC software. Gene-level read counts were computed using HT-Seq-count (Anders *et al.* 2015) with ‘union’ mode and genes with less than 10 aligned reads across all samples were discarded prior to analysis leading to 15,155 genes. Differential gene expression analysis between WKY and LEW BMDMs was performed using DESeq2 (Love *et al.* 2014) and significantly differentially expressed genes were reported at the 5% FDR level. The visualizations of the expression levels with gene structure were created with DEXSeq (Anders *et al.* 2012).

ChIP-seq data from BMDM isolated from the WKY and WKY.L*Crgn2* congenic strains (in which the LEW Crgn2 QTL was introgressed onto the WKY background) were retrieved from (Hull *et al.* 2013; Srivastava *et al.* 2013) and re-analyzed with respect to the *Ifitm3* locus. This congenic model (WKY.L*Crgn2*) has been extensively studied in previous studies where it has been shown that JunD expression levels are significantly higher in WKY when compared with the congenic (Hull *et al.* 2013) and that the canonical binding of AP-1 is significantly greater in WKY compared to WKY.L*Crgn2* (Behmoaras *et al.* 2008). Briefly, ChIP was performed with a JunD antibody (Santa Cruz sc74-X) and a negative IgG control (sc-2026). Single read library preparation and high throughput single read sequencing for 36 cycles was carried out on an Illumina Genome Analyser IIx and sequencing of the ChIP-seq libraries was carried out on the high throughput Illumina Genome Analyzer II. Initial data processing was performed using Illumina Real Time Analysis (RTA) v1.6.32 software (equivalent to Illumina Consensus Assessment of Sequence and Variation, CASAVA 1.6) using default settings. Quality filtered reads were then realigned to the rn4 using the Burrows Wheeler Alignment tool v0.5.9 (BWA). Read ends were trimmed if Phred-scaled base quality scores dropped below 20. For the ChIP-seq analysis presented in Figure 3g, differences in JunD binding were assessed only within a 700bp region spanning the *Ifitm3* gene promoter, which included the 600 bp-long DMR identified by ABBA at this locus. ChIP-seq differences were assessed by means of Fisher’s exact test on the ChIP-seq counts (normalized for library size) in WKY L*Crgn2* and LEW strains, respectively, using a sliding window of 50bp. This locus-specific analysis identified a single 50bp window with differential JunD binding with FET p-value<0.05 that overlapped with JunD TFBS motifs identified by HOMER (see above).

### Software and data availability

ABBA is implemented as a Perl/R program, which is available with instructions for download at *abba.systems-genetics.net* or via http://www.mrc-bsu.cam.ac.uk/software/bioinformatics-and-statistical-genomics/. The data is available on Gene Expression Omnibus (GEO), <u>https://www.ncbi.nlm.nih.gov/geo/</u>, under the accession number GSE84719.

## RESULTS

We employ a fully Bayesian approach (a Bayesian structured generalized mixed additive model with a latent Gaussian field) which models the random sampling process of the WGBS experiment (the number of methylated/unmethylated reads distributed as non-Gaussian response variable) and where all the unknown quantities are specified by probability distributions. To perform inference ABBA takes advantage of the Integrated Nested Laplace Approximation (INLA) (Rue *et al.* 2009), a new inferential tool for latent Gaussian models. INLA provides approximations to the posterior distribution of the unknowns. These approximations are both very accurate and extremely fast to compute compared to established exact sampling-based methods such as Markov chain Monte Carlo (Gilks *et al.* 1996) (MCMC) or Sequential Monte Carlo (Doucet *et al.* 2001) (SMC). Our new proposed algorithm ABBA is therefore the combination of an <u>approximate</u> inferential procedure with a fully <u>Bayesian</u> model tailored for <u>bisulphite</u> sequencing <u>analysis</u>.

ABBA calculates the posterior methylation probability (PMP) at each CpG site based on an estimate of the posterior probability of a smoothed unobserved methylation profile. It also identifies DMRs at a specified FDR by contrasting PMPs across the whole-genome between two groups, e.g. cases and controls. Several intrinsic features of WGBS data are incorporated into ABBA: for instance, the variability in DNA methylation between the (experimental) replicates within each group is modeled through a random effect with a specific within-group variance (**Fig. 1a**). The correlation of DNA methylation patterns is encoded in the latent Gaussian field equation, which reflects the neighborhood structure of the model and automatically adapts to the changes in the underlying data. In particular, the *a priori* correlation between neighbouring CpGs’ methylation profiles depends on the distance between them, as it decreases as this distance increases (**Fig. 1b**). Rather than relying on a user-defined value to parameterize it (e.g., kernel bandwidth or window size) or fixing it by an automatic procedure (for instance through an empirical Bayes approach), ABBA assigns a prior distribution on the parameters of the latent Gaussian field equation, thus fully accounting for the uncertainty about these quantities. This specification is key in our model since the data-adaptivity of the degree of smoothing conforms better to the data than assuming fixed values. All these features allow our model to adjust routinely to real-world scenarios, providing an automatic way to describe the WGBS data without requiring any user-defined parameters (Yu and Sun 2016b). Full technical details of ABBA algorithm can be found in the Materials and Methods.

We benchmarked ABBA and compared it against recently proposed methods (MethylKit (Akalin *et al.* 2012), MethylSig (Park *et al.* 2014), DSS/DSS-single (Feng *et al.* 2014; Wu *et al.* 2015), simply DSS hereafter, BSmooth (Hansen *et al.* 2012), metilene (Jühling *et al.* 2015) and the univariate Fisher’s exact test (FET)). All methods were run using their default parameterization and for the FET we pooled data from different replicates. To ensure a fair comparison, we used WGBSSuite (Rackham *et al.* 2015) to generate a large number of diverse datasets that were independent of the underlying statistical models of ABBA and of the other methods. Briefly, we simulated *in-silico* datasets to assess the performance of each method under several scenarios, which reflect differences in data integrity and quality of the signal that can occur as a result of biological and experimental phenomenon. The parameters considered were the following: the number of replicates within each group (*r*), the average read depth per CpG, the level of noise variance (*s*_0_), the methylation probability difference between the two groups (*Δmeth*) and the switching of methylation status of CpG sites between the two groups (*δ*) (see **Methods** for details). For each simulated case we generate five replicates and we compared the accuracy of the CpGs called as being contained within DMRs by each technique with the true simulated DMRs. To quantitatively assess the performance of ABBA with respect to competing methods, we evaluated false-positive and false-negative rates of CpG sites and generated receiver operator characteristic (ROC) curves. We focus on the partial area under the ROC curve (or pAUC) at a specificity of 0.75. The pAUC is considered to be more practically relevant than the area under the entire ROC curve (Ma *et al.* 2013) since in typical genomics studies only the features identified at very low false positive rates are selected for further biological validation.

All results of the benchmark are detailed in **Supplementary Figures 5-7**. In **Fig 2a** we show representative ROC curves from a specific combination of parameters whilst in **Fig. 2b** we summarize the performance over all combinations of parameters by displaying the best performing method based on its pAUC. Specifically, in **Fig. 2b** the color code in the “benchmark grid” indicates the best performing method for each of the 216 simulated scenarios. For instance, in **Fig. 2a** the top left panel (i) shows the ROC curves for all methods considered under a simulated dataset with *s*_0_ = 0.1, *Δmeth* = 30%, *r* = 1, average read depth per CpG of 10x and *δ* = 0. For this combination of parameters we compared the pAUC of each approach, which shows that ABBA is the best performing method. Accordingly, in **Fig. 2b** the square in the grid that represents this parameter set (indicated by (i) in the figure) is coloured black (ABBA). Examples of other ROC curves for specific combinations of parameters are reported in **Fig. 2a** (i-vi) and the corresponding best performing methods are indicated in **Fig. 2b**. In some simulated cases (e.g., with high levels of *δ* = 10%) the ROC curves and corresponding pAUC do not distinguish unambiguously the best performing method (e.g., **Fig. 2a** – panel (vi)). In these cases when the pAUC of two methods are very similar (±1%) we report the colours of both methods, e.g., black and red colours in the same square to indicate similar performance of ABBA and DSS (**Fig. 2b**). For the metilene approach (Jühling *et al.* 2015) (which was run using its default parametrization) we noticed that ROC curve analysis was not suitable to compare its perfomance with other methods. Specifically, for metilene we found that it was not possible to assess both specificity and sensitivity across the wide range of DMRs and scenarios simulated in our study. Representative examples for the ROC curves obtained by running metilene (and other approaches) on the simulated data are provided in **Fig. 2a** and in **Supplementary Figure 8**.

**Figure 2.**
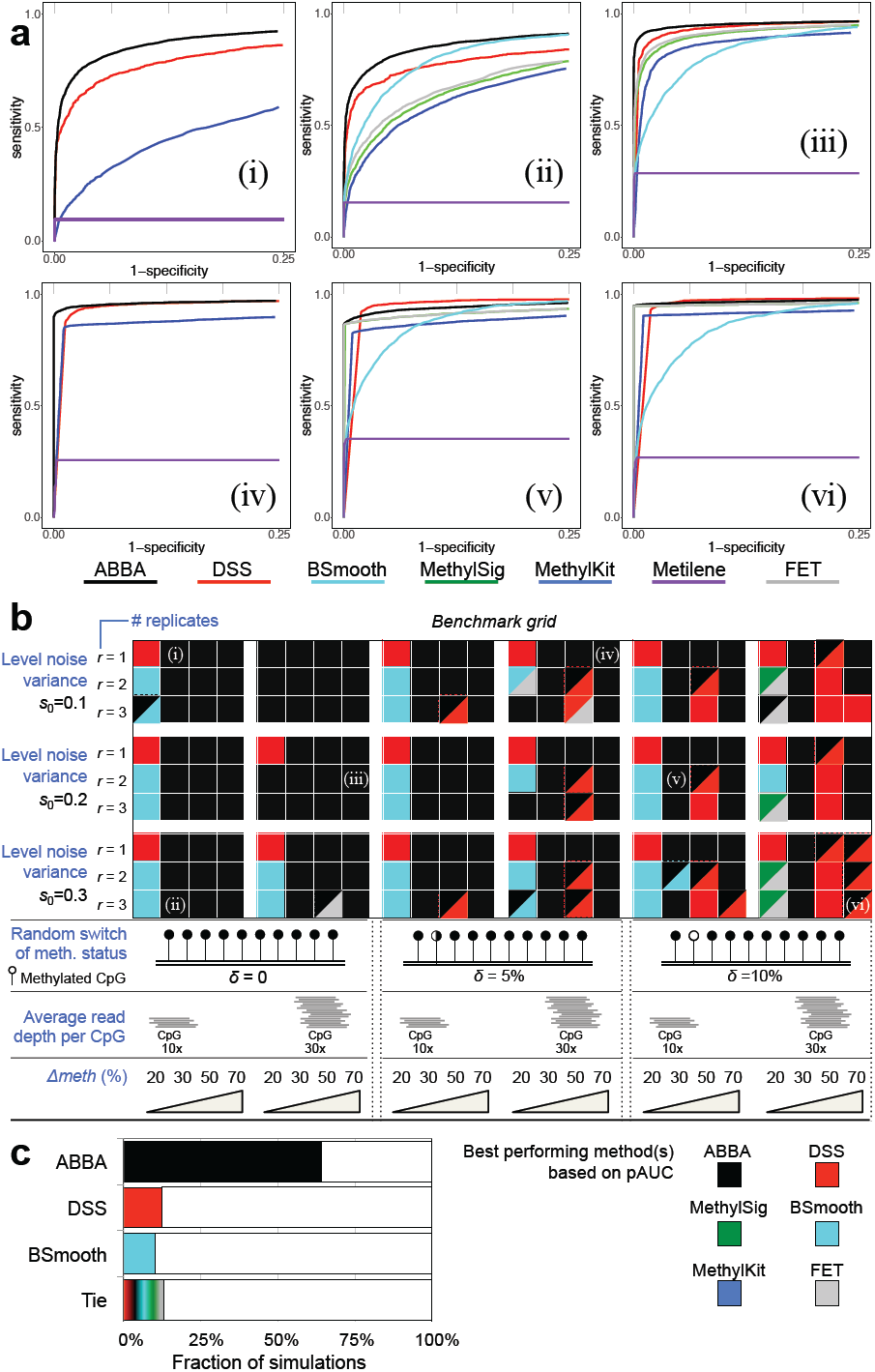
Benchmarking results. (**a**) ROC curves for selected combinations of parameters: (i) *s*_*0*_ = 0.1, *Δmeth* = 30%, *r* = 1, average read depth per CpG of 10x, *δ* = 0; (ii) *s*_*0*_ = 0.3, *Δmeth* = 30%, *r* = 3, average read depth per CpG of 10x, *δ* = 0; (iii) *s*_*0*_ = 0.2, *Δmeth* = 70%, *r* = 2, average read depth per CpG of 30x, *δ* = 0; (iv) *s*_*0*_ = 0.1, *Δmeth* = 70%, *r* = 1, average read depth per CpG of 30x, *δ* = 5%; (v) *s*_*0*_ = 0.2, *Δmeth* = 30%, *r*=2, average read depth per CpG of 10x, *δ* = 10%; (vi) *s*_*0*_ = 0.3, *Δmeth* = 70%, *r* = 3, average read depth per CpG of 30x, *δ* = 10%. For each of this combination of parameters, the corresponding best method based on its pAUC is indicated in the benchmark grid below. In (i) and (iv) ROC curves are reported only for the methods that can analyze WGBS data generated from one biological sample. (**b**) Global snapshot of the method’s performance across 216 simulated datasets. A given combination of parameters is indicated by a square in the benchmark grid, and for each square we calculated the pAUC for each method and determined which method had the overall best pAUC (i.e., pAUC_method_1_ > pAUC_method_2_). Colours in the benchmark grid indicate which method had the best performance. When pAUC of two methods are similar (±1%) we report the colours of both methods (e.g., black and red colours in the same square indicate similar performance of ABBA and DSS). The six selected combination of parameters for which the ROC curves are reported in panel (a) are indicated within the benchmark grid: (i, ii, iii, iv, v and vi). All ROC curves are reported in **Supplementary Figures 5-7**. (**c**) For the three best performing methods (ABBA, DSS and BSmooth) we report the percentage of simulated scenarios in which each method resulted to be the best based on the pAUC comparison. “Tie” indicates the proportion of simulated scenarios in which the pAUCs of any two methods were similar (i.e., pAUCs ±1%) and it was not possible to single out a single best performing approach.

Considering all 216 simulated datasets and comparing the pAUCs obtained by each approach across all combinations of parameters, ABBA (black) showed to be the best performing method in 139 (64%) cases (**Fig. 2b-c**). The two other competitive methods were DSS and BSmooth, which show to be the best performing approach only in 26 (12%) and 22 (10%) simulated cases, respectively (**Fig. 2b-c**). In 28 (13%) cases different methods showed very similar performance (i.e., pAUCs ±1%), and in 17 simulations ABBA and DSS showed to have comparable performance. Looking at the detailed ROC curves reported in **Supplementary Figures 5-7**, we notice that while ABBA was the best method across all simulations (**Fig. 2c**), its performance diminished for simulated datasets with very small methylation probability difference between the two groups. In particular, for most of the simulated scenarios with *Δmeth* = 20%, BSmooth showed very good and robust performance, while DSS was consistently the best performing method when *r* = 1 and *Δmeth* = 20%, **Fig. 2b**. However, we highlight that such small differences in DNA methylation (i.e., *Δmeth* ≤ 20%) are unlikely to have an important biological effect, and the most commonly observed effect sizes for DMR range between 20 and 40%, as previously reported (Ziller *et al.* 2015). In the range *Δmeth* ≥ 30%, ABBA was the best performing method in 132 (81%) simulations, while DSS was the best performing method only in 10 (6%) simulated cases and, notably, BSmooth was never the best single performing method (BSmooth showed similar performance of ABBA in only one simulated case) (**Fig. 2b**).

Specific observations have to be addressed when high levels of errors due to the switching of methylation status of CpG sites between the two groups have been simulated. In these scenarios, it was more difficult to single out a method that outperforms all competing approaches. However, when *δ* was as high as 10% (i.e., 1 in 10 CpGs is misclassified as unmethylated or vice versa), we observed that ABBA was the best single method in 33 (46%) of 72 simulated scenarios, whereas DSS and BSmooth performed as the best method in 16 (22%) and 7 (10%) of cases, respectively, and in other 10 cases ABBA and DSS have comparable performance. The latter was more apparent when large probability differences between the two groups were simulated (*Δmeth* = 50% or 70%).

We then explored whether non-homogeneous, spatially correlated read depth has an effect on ABBA’s performance. In order to capture spatially correlated read depth from real data we sampled 5,000 contiguous CpGs from WGBS data (generated in rat macrophages, see below and **Methods** for details) and then varied other parameters (*r* and *Δmeth*) using WGBSSuite as described above. In these “data-derived” simulated datasets the read depth was correlated with the distance between CpGs (**Supplementary Figure 9a**). The results of the benchmark using read depth taken from real data were very similar to those obtained using read depth simulated by means of a Poisson distribution (see **Methods**). Regardless of whether “data-derived” or “Poisson-simulated” read depth was used in our simulations, ABBA was the best performing method to recall DMRs (representative examples are reported in **Supplementary Figure 9b**). While heterogeneous levels of read depth impact on the single base probability of methylation, the hierarchical model underlying ABBA borrows information across the sequence analyzed, it turns out that ABBA posterior estimates are less sensitive to different levels of the read depth.

Taken together our simulation study shows that while individual approaches can be very powerful in detecting DMRs under specific scenarios (notably, DSS with *r* = 1 and BSmooth with *Δmeth* = 20%), their performance can vary (and drop) significantly for different choices of the parameters tested in our simulations (at least within the parameter-space considered here). In contrast, we show that, on the whole, ABBA is the best performing method across a large number of parameters’ combination tested and accurately identifies DMRs in the large majority of simulated cases (**Fig. 2c**). Specifically, ABBA’s performance was the highest in the detection of biologically meaningful changes in DNA methylation (*Δmeth* ≥ 30%) and when little or no errors due to random switching of methylation status of CpG sites between the two groups are present in the data.

DNA methylation is emerging as a major contributing factor in several human disorders (Zoghbi and Beaudet 2016), including important autoimmune diseases like systemic lupus erythematosus (SLE) (Wu *et al.* 2016). For instance, differential DNA methylation analysis in CD4+ T cells in lupus patients compared to normal healthy controls identified several genes with known involvement in autoimmunity (Jeffries *et al.* 2011). Here, to illustrate the practical utility of ABBA for differential methylation analysis in disease, we generated WGBS data in an established experimental rat model of crescentic glomerulonephritis (CRGN)(Aitman *et al.* 2006). In this model, we and others have previously shown that susceptibility to CRGN is mediated by macrophages (Behmoaras *et al.* 2008; Page *et al.* 2012); therefore, we assayed CpG methylation at single-nucleotide resolution by WGBS in primary macrophages derived from Wistar Kyoto (WKY) and Lewis (LEW) isogenic rats (two strains discordant for their predisposition to develop CRGN). We used ABBA to carry out genome-wide differential DNA methylation analysis in primary bone-marrow derived macrophages (BMDM) derived from the disease-prone rat strain (WKY, *r* = 4) and control strain (LEW, *r* = 4) - see **Methods** for additional details on WGBS data generation and processing. Briefly, in our ABBA analysis of the macrophage methylome, we used the following (default) settings: a minimum of 5 CpG and at least 33% difference in DNA methylation between the disease and control macrophages to identify DMRs. This choice was motivated and supported by data on the local topology of CpG sites in the methylome showing the vast majority of the CpG clusters are in the range of 1–11 CpGs (Lövkvist *et al.* 2016) and to increase true positive rate in our DM analysis, following previous assessment and recommendations for methylation analysis using WGBS data (Ziller *et al.* 2015).

Using an FDR cutoff of 5%, ABBA identified 1,004 DMRs genome-wide, with 1.07% falling within an annotated CpGI and 6.78% within an annotated CpGS (**Fig. 3a**). For comparative purposes we also used DSS (since this method performed very similarly to ABBA in several simulated cases, **Fig. 2**) to identify DMRs genome-wide, which resulted in only 207 regions with significant differential methylation (uncorrected p-value threshold = 10-3, using the default parameters of DSS). Of the 1,004 DMRs identified by ABBA, 427 overlapped with annotated genes (**Supplementary Table 3**), and there was a significant enrichment for DMRs occurring within 1kb of the gene boundaries (p-value<0.001), within exons (p-value<0.05) and introns (p-value<0.05), **Fig. 3b**. The genes that are within 1kb of a DMR were enriched for pathways relevant to the pathophysiology of CRGN, including MAPK signalling (Ryan *et al.* 2011), Phosphatidylinositol signalling (Wu *et al.* 2014) and Fc gamma R-mediated phagocytosis (Page *et al.* 2012) (**Fig. 3c**). For comparison, the 207 DMRs identified by DSS overlapped with 45 genes (**Supplementary Table 4**), which were enriched only for RNA degradation and metabolic pathways. The analysis of real WGBS data by DSS highlighted how the choice of parameters (in this case related to the size of the moving average window in the smoothing procedure) can affect the results. Since the window size in DSS is a user-defined parameter, we performed the analysis with DSS using three different windows (50 bp, 100 bp, 1,000 bp) in addition to the default window size of 500 bp. Each of the four window sizes identified a different number of DMRs, which overlap with different genes (**Supplementary Figure 10a**) and have varying distributions of DMR lengths (**Supplementary Figure 10b-e**). The genes identified by DSS when a window of 50 bp is used showed no significant enrichment for pathways, while the results obtained for 100 bp and 1,000 bp windows showed a significant enrichment for RNA degradation. These analyses highlight how the arbitrary choice of parameters related to the degree of smoothing can affect greatly the results of a genome-wide DM analysis as well as the downstream annotation of the genes overlapping with DMRs. In contrast, ABBA automatically adapts to different correlation structures in DNA methylation levels across the genome without requiring any user-defined parameters related to the smoothing procedure.

**Figure 3.**
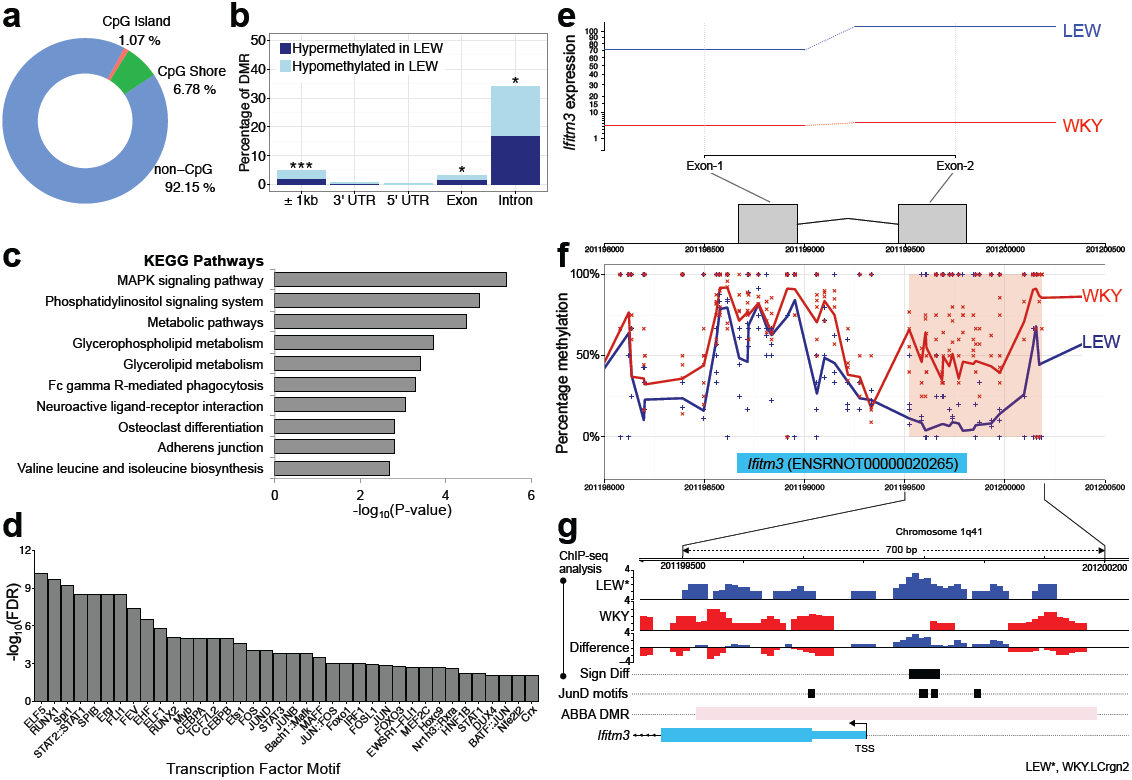
ABBA analysis of WGBS in rat macrophages. (**a**) CpG-based annotation 1,004 DMR between WKY and LEW macrophages showing significantly higher proportions of CpGI and CpGS than those that would be expected by chance (p-value<0.009 for CpGI and p-value<0.001 for CpGS, respectively, obtained by 1,000 randomly sampled datasets of 1,004 CpG-matched regions). (**b**) Proportions of DMRs in different genomic features of overlapping genes. Feature annotation was retrieved from UCSC genome browser (RN4). (c) KEGG pathway enrichment for the genes overlapping with DMRs. Only significant pathways are reported (FDR<1%). (**d**) Enrichment for the TFBS within the DMRs was when compared to CG matched regions of the genome (FDR<0.05). (**e**) RNA-seq analysis in WKY and LEW macrophages shows lack of *Ifitm3* expression in WKY rats. (**f**) Percentage methylation at each CpG in WKY (crosses) and LEW (plus) and smoothed average methylation profiles by ABBA. The pink box highlights the significant DMR identified by ABBA (FDR<5%). (**g**) ChIP-seq analysis for JunD in LEW.LCrgn2 (LEW*) and WKY macrophages identified a single region with differential binding of JunD (p-value<0.05, Sign Diff row, black box). Units on the y-axis refer to relative ChIP-seq coverage with respect to the control. This region overlapped with two (out of four) JunD binding sites motifs identified within the gene promoter (±500bp around the TSS). ABBA DMR, differentially methylated region identified by ABBA. TSS, transcription start site. *, p-value<0.05, ***, p-value<0.001

As DNA methylation can affect gene expression by interfering with transcription factor binding, we performed a transcription factor binding site (TFBS) analysis of the DMRs (**Fig. 3d**). This revealed significant enrichment for several TFs, including the ETS transcription factors family and a number of proteins that make the AP-1 TF complex (JUNB, FOS, JUN and JUND), which have been previously linked with CRGN (Behmoaras *et al.* 2008),(Raffetseder *et al.* 2004). To further investigate the potential effect of the changes in DNA methylation identified by ABBA, we carried out differential expression (DE) analysis in macrophages from WKY and LEW rats by RNA-seq (see **Methods** for details). The list of DE genes (n=910, Benjamini–Hochberg (BH)-corrected p-value<0.05) was crosschecked with the genes impacted by DMRs (above), identifying 48 genes with both significant differential methylation and differential expression (**Supplementary Table 5**). We observed the “textbook” model describing DNA methylation regulating transcription via the promoter region (i.e., hypermethylation in the promoter associated with transcriptional repression, see below) as well as widespread methylation changes in the genes body and 3’UTR associated with both gene repression and activation. The genes with concordant promoter hypermethylation and transcriptional repression, *Ifitm3*, *Ydjc* and *Cd300Ig* were investigated in more detail since the gene’s promoter is a key regulatory region where the effect of DNA methylation is more clearly understood. We found the biggest change in mRNA expression was in interferon induced transmembrane protein 3 (*Ifitm3)*, with mRNA from this gene being almost undetected in unstimulated WKY macrophages (**Fig. 3e**). This observation is consistent with the differential methylation status of the promoter of *Ifitm3*, where the WKY rats had higher levels of methylation than the LEW rats (**Fig. 3f**). To further support the identification of differential methylation at the *Ifitm3* gene we checked whether other methods identified the same DMR. While MethylSig failed to identify significant DMR and BSmooth identified a large and unspecific genomic area as differentially methylated, DSS provides highly consistent results with ABBA, identifying differential methylation at the same region at the *Ifitm3* gene promoter (**Supplementary Figure 11**).

We have previously shown that JunD (AP-1) transcription factor is a major determinant of CRGN in WKY rats (Behmoaras *et al.* 2008) and others have shown that AP-1 is methylation sensitive (Ogawa *et al.* 2014). Therefore we scanned the DMR (spanning 600 bp) for canonical JunD binding site motifs, and identified three putative regions in the promoter region of *Ifitm3* (**Fig. 3g**). In addition, we re-analyzed ChIP-seq data for JunD transcription factor in BMDM derived from WKY and a congenic strain from LEW (see **Methods** for details). This analysis identified significant differences in JunD binding between WKY and LEW-congenic strain that overlapped with two of the four TFBS identified at the *Ifitm3* promoter (**Fig. 3g**). The combined evidence provided by our ABBA analysis and RNA-seq/ChIP-seq data therefore suggests that the effect of DNA methylation of the *Ifitm3* gene promoter in WKY rats (prone to develop CRGN) may be restricting the binding of transcription factors such as JunD and, as a consequence, the gene is almost not expressed (<1 TPM) in unstimulated macrophages of WKY rats.

## DISCUSSION

As the cost of genome sequencing technologies continues to drop, it will soon become commonplace to perform comprehensive methylome analyses, using WGBS or other high-throughput techniques that allow the unbiased genome-wide quantification of DNA methylation at a single base-pair resolution. However, high-resolution data generation is only the first step towards the identification of genomic loci and eventually genes with altered methylation levels associated with a given disease, phenotype or developmental stage. The number of DNA methylation datasets available in the public domain is expected to grow; therefore, it becomes necessary to provide the scientific community with analytical tools for a reliable and reproducible identification of differential methylation, and facilitate large epigenome-mapping projects and epigenome-wide association studies (Bock 2012).

Beyond statistical power considerations specifically related to the sample size (Rakyan *et al.* 2011) or interpretability of epigenome-wide association studies (Birney *et al.* 2016), our ability to identify accurately changes in DNA methylation localized to specific genomic loci (genes) is also influenced by multiple factors inherently correlated to data quality. These include the within-group heterogeneity, the level of noise, the presence of known genetic covariates (Zhang, 2015) and non-genetic confounding factors (e.g., batch effects) as well as features such as sequencing depth (Ziller *et al.* 2015) or errors due incomplete bisulphite conversion (Genereux *et al.* 2008). Therefore, any analytical tool that can account for all these factors will reduce the number of false positives maximizing the sensitivity and call the regions of interest (i.e., differentially methylated) as accurately as possible. With this in mind, we designed a differential methylation analysis tool (ABBA) that is robust to different experimental and technical variables (see **Fig. 2**), and that adapts automatically to the varying genomic context and local topology of CpG sites affecting methylation levels. In particular, the automatic adaptation to different correlation structures in CpG methylation levels (without requiring user-defined parameters about the degree of smoothing) as well as the ability of modelling its decay as the function of the genomic distances between CpGs allow ABBA to adapt routinely to methylation changes that occurs with different scales and non-uniform rates across the genome. The importance of the genomic context in the methylome and the local topology of CpG sites have been recently investigated, showing, amongst other features, that methylation at small CpG clusters is more likely to induce stable changes in DNA methylation (Lövkvist *et al.* 2016).

From a user’s perspective, ABBA treats WGBS-seq data in a general way with no specification of parameters related to the level of data smoothing (such as window size or kernel bandwidth), thus allowing for a great deal of automation. This also facilitates the WGBS analysis when the values of the parameter settings (that may largely affect the accuracy of DM identification) are not known. Our fully Bayesian approach can be also easily modified to include covariates and non-genetic confounding factors through random effects, beyond the replicates level. It also allows the specification of covariates that are informative about the methylation profiles by adding prior biological information to the linear predictor *μ*_*ig*_ in (6). While these alterations can be done in our model with a simple modification of the code and with negligible further computational costs, non-parametric smoothing techniques (spline-(Hansen *et al.* 2012), kernel (Hebestreit *et al.* 2013)- and moving average-based smoothing (Feng *et al.* 2014)) do not possess the same straightforward flexibility nor alternative approaches based on Hidden Markov Models (Yu and Sun 2016a), (Kuan and Chiang 2012), (Sun and Yu 2016).

Our extensive simulation studies (**Fig. 2**) and differential DNA methylation analysis in glomerulonephritis (**Fig. 3**) showed that ABBA is a powerful approach for the identification of DMRs from WGBS single-base pair resolution methylation data. While individual methods such as BSmooth (Hansen *et al.* 2012) or DSS (Feng *et al.* 2014; Wu *et al.* 2015) showed a very good power to detect DMRs under specific scenarios and conditions, ABBA retained a high degree of robustness of the results with respect to a wider range of factors (parameters) affecting WGBS data integrity and quality, including sequencing coverage, number of replicates or different noise structures. This is particularly appealing in cases when considerable efforts have been expended toward generation of large-scale WGBS data from heterogeneous systems, e.g., the ENCODE project (Bernstein *et al.* 2012), and data quality can vary across experimental conditions and laboratories. As proof of concept of ABBA’s application to real data analysis, we used an established experimental model system of glomerulonephritis (Aitman *et al.* 2006) to identify changes in DNA methylation associated with disease. In this, we employed ABBA to analyze ~15 million CpG sites genome-wide in primary bone-marrow derived macrophages derived from WKY and LEW rats and identified >1,000 significant DMRs at 5% FDR level. A comparative analysis using DSS (the most competitive approach from our simulation study) did not provide the same level of biological insight both in terms of significant pathway enrichments and in robustly identifying DMRs across user-defined parameters. To highlight this point, we showed how the results of DSS were greatly affected by the choice of the window size.

Furthermore, we have shown how integrating the DMR results provided by ABBA with other ‘omics’ data (RNA-seq and ChIP-seq generated in the same experimental system), enabled us to generate new hypotheses for the mechanism underpinning the disease, revealing a candidate gene (*Ifitm3*) for the susceptibility to glomerulonephritis. These findings on *Ifitm3* in rat glomerulonephritis merit further discussion. *Ifitim3* has a known role in viral resistance, a central part of innate immunity, and is inducible by both interferon (IFN) types I and II (Everitt *et al.* 2012). Notably, type II IFN signaling has been implicated in the pathogenesis of nephrotoxic nephritis and other “planted” antigen models of CRGN (Kitching *et al.* 2004), although DNA methylation has not previously been examined in this context. With regards to type I IFN, recent genome-wide DNA methylation analysis of T-cells, B-cells and monocytes has shown that patients with SLE, a frequent autoimmune cause of CRGN, have severe hypomethylation near to genes involved in type I IFN signaling (Absher *et al.* 2013). In addition, DNA methylation alterations in IFN-related genes, including *Ifitm3*, have been previously observed and proposed to contribute to the pathogenesis of other autoimmune diseases such as primary Sjögren’s syndrome (Gottenberg *et al.* 2006). Regarding the role of *Ifitm3* gene, it has been shown to directly interact *in vivo* and *in vitro*, with osteopontin, a matricellular protein, whose transcription is mediated by the AP-1 TF family (El-Tanani *et al.* 2010). Furthermore, osteopontin has been also previously associated with SLE (Rullo *et al.* 2013) and ANCA-associated vasculitis (Lorenzen *et al.* 2010) another frequent cause of CRGN. Therefore, our ABBA analysis of WGBS data in primary macrophages from a rat model of CRGN allowed us to propose an AP-1-mediated role for *Ifitm3* in glomerulonephritis. While a role for IFN-signaling genes in autoimmune disease has been previously suggested, our findings on methylation alteration of the *Ifitm3* gene associated with glomerulonephritis in the rat might suggest future directions for the study of the pathogenesis and to develop treatments of CRGN.

In a wider context, the role of methylation is dependent on the location with respect to the gene body and regulation functions. Methylation in a CpGI-depleted promoter, such as the promoter region of *Ifitm3* gene (according to UCSC genome browser (RN4)), is associated with repression that maybe due to interference with transcription factor binding. Conversely, methylation in the gene body is positively associated with active transcription as methylation can be caused by transcriptional elongation (Schübeler 2015). Methylation within a gene body can also act as an insulator for repetitive and transposable elements or distal intronic enhancers, on which the methylation would have no regulatory effect on the gene in which it resides (Jones 2012). Given the complexity of these regulatory functions of methylation, the ability of our approach to accurately identify changes in DNA methylation that are localized to specific regions is likely to facilitate our understanding of the complex relationships between methylation and gene regulation. As exemplified by our integrative analysis of the of the *Ifitm3* locus, we anticipate that the ABBA results for differential DNA methylation should be integrated with additional transcriptional and epigenetic data in order to better define hypotheses on specific regulatory mechanisms.

In summary, we show how ABBA provides a flexible and user-friendly automatic framework for the identification of differential methylation that is robust across a wide range of experimental parameters, an approach that we have also applied to identify changes in macrophage DNA methylation in glomerulonephritis.

## ACKNOWLEDGEMENTS

The authors are thankful to the two anonymous referees whose meticulous attention to their refereeing task has resulted in substantial improvements in our presentation.

## FUNDING

This research was funded by Engineering and Physical Sciences Research Council Grant EP/K030760/1 (L.B.), The Alan Turing Institute under the EPSRC grant EP/N510129/1 (L.B., P.D.), Royal Society IE110977 (L.B., P.D.), European Union (European Social Fund - ESF), Greek national funds through the Operational Program "Education and Lifelong Learning" of the National Strategic Reference Framework (NSRF), project ARISTEIA (P.D.), Duke-NUS Medical School and Singapore Ministry of Health (O.J.L.R., E.P.), a Medical Research Council Chain-Florey fellowship (T.O.), the Medical Research Council (MR/M004716/1 to J.B. and E.P.) and by Kidney Research UK - RP9/2013 (J.B.). The funders had no role in study design, data collection and analysis, decision to publish, or preparation of the manuscript.

## AUTHOR CONTRIBUTIONS

L.B. and E.P. initiated, directed and supervised the project. P.D. and L.B. conceived the statistical model and the computational approach. P.D., E.V. and L.B. wrote the initial algorithm that was further developed by O.J.L.R. and L.B. to the presented approach. T.O. and E.P. generated WGBS data in the rat. S.R.L., O.J.L.R. and E.P. carried out analysis of WGBS and RNA-seq data in the rat and interpreted the results. N.H. and P.K.S. carried out ChIP-seq and TFBS analyses. J.B. provided RNA-seq and ChIP-seq data in the rat. O.J.L.R., L.B. and E.P. wrote the manuscript with input from all authors. All of the authors read and approved the final manuscript.

